# Molecular basis for the regulation of retroviral nucleosomal integration by chromatin compaction

**DOI:** 10.1101/2025.05.05.652197

**Authors:** D. Lapaillerie, Y. Zgadzay, F. G. Martins, C. Bertinetti, M. Autin, J. Batisse, C. Tumiotto, P. Lesbats, M. Ruff, S. F. Sousa, V. Parissi

**Author notes:** Dynamic of the interactions between viral and cellular chromatins CNRS GDR 2194 (DyNAVir).

## Abstract

Cellular chromatin represents the first nonreversible contact point between the genomes of incoming infectious agents, such as integrative viruses, and their hosts. The integration of retroviral genomes requires a functional association between the viral integration complex (intasome) and host chromatin, mediated through multiple interfaces between the integrase, target DNA, and histone components of the nucleosome. Previous studies have shown that these associations are regulated by cellular factors, such as LEDGF/p75 for lentiviruses, and by the degree of compaction of the chromatin surrounding the insertion site. However, the molecular mechanisms underlying the regulation of access to local nucleosomal functional interfaces remain elusive. In this study, we dissected how incoming intasomes engage nucleosome surface to achieve efficient integration. Combining biochemical approaches with molecular docking analysis, we demonstrated that HIV-1 and PFV intasomes distinctly and specifically interact with nucleosome surfaces. Mapping these interfaces onto a compacted trinucleosomal structure and simulating the dynamic docking of the intasome at these sites revealed how neighboring nucleosomes modulate the functional binding of the HIV-1 intasome to the catalytic target nucleosome by masking these functional interfaces. In contrast, the lower susceptibility of the PFV intasome to chromatin compaction was due to the persisting accessibility of active nucleosomal interfaces. Together, these data provide the first molecular and structural insights into how chromatin compaction influence retroviral integration. Our results especially show how nucleosome-intasomes docking sites participate in modulating the sensitivity of the retroviral integration to chromatin structure. Overall, our data reveal that HIV-1 and PFV integration rely on nucleosomes with distinct structural and functional properties at the insertion site to form an active strand transfer complex. This work further demonstrated that retroviruses have evolved distinct strategies to engage suitable chromatin structures for efficient integration highlighting a divergence in retroviral adaptation mechanisms.

## INTRODUCTION

Retroviral infection requires stable integration of viral DNA into the host chromosomes. In infected cells, integration is carried out by the pre-integration complex (PIC), which contains viral DNA bound to the virally encoded integrase (IN) enzyme, forming the intasome, and other viral/host factors whose nature remains poorly known. Integration occurs selectively into specific “hot spots” of host chromosomes (1). Many parameters govern this insertion site preference including nuclear import processes, cellular targeting pathways and nucleus architecture (2). Many studies also indicate that structural features of chromatin such as targeting as epigenetic marks, nucleosomal compaction and remodeling activities (extensive literature), are important for integration. However, to date, the molecular mechanisms that guide the PIC and enable the insertion of PIC-associated viral DNA into specific hot spots of human chromosomes are not fully understood. Among these factors, the final interaction of the intasome with the nucleosomes forming the target capture complex (TCC) and finally the strand transfer complex (STC) plays a central role in the targeting process. The chronology of these events as well as the determinants for this final chromatin interaction remain to be fully understood.

Within the chromatin, the host target DNA is compacted as a mixture of nucleic acids and proteins found in the nucleus of eukaryotic cells that help to condense and organize genetic information. The nucleosome represents the core repeating unit of chromatin and constitutes 147 bp of DNA wrapped around a histone octamer (3). Nucleosomes are dynamic structures that can be assembled and disassembled by chaperones or chromatin remodelers and modified through posttranslational modifications. The local environment around the nucleosome imparted by these factors is thus expected to play an important role in establishing a successful integration event. *In vitro*, integration into nucleosomes can be recapitulated by the assembly of intasomes formed by multimers of IN, the number of which varies from 4 to 16 depending on the virus (4), and viral DNA. The use of these reconstituted intasomes in *in vitro* integration assays highlighted differences between retroviruses, showing their distinct preferences for chromatin compaction, which was further confirmed *in cellulo* (5, 6). Functional association with nucleosomal DNA requires the accessibility of the intasome/nucleosome interfaces, which may define the preference of the different intasomes for specific chromatin compaction levels and structures. Previously, direct interactions between retroviral intasomes and both the DNA and the histone components of the nucleosome were reported. Indeed, while the PFV intasome binds the H2A/H2B dimer interfaces (7), HIV-1 IN was reported to interact preferentially with histone H4 *in vitro*, leading to structural changes within the IN and stimulating integration (8, 9). A distinct association with chromatin was also detected in the context of human chromosomes via *in vitro* direct chromatin binding assays highlighting the importance of neighboring chromatin in regulating these functional associations (10).

Taken together, these data suggest that PFV and HIV-1 intasomes employ distinct ways to engage nucleosomes. In this process, various intasomes may interact differentially with nucleosomes, exhibiting distinct behaviors at chromatin contact sites. Integration sensitivity toward the chromatin structure neighboring the insertion site may rely on these different interfaces. In this work, we determined the influence of intasome/nucleosome interfaces on chromatin structure sensitivity and better elucidate the mechanistic basis of the selection of specific regions of chromatin for integration. For this purpose, we compared HIV-1 and PFV retroviral intasomes, which clearly show clear distinct chromatin preferences, using *in vitro* integration assays and biochemical approaches with either isolated nucleosomes or trinucleosomes, allowing us to analyze the impact of direct neighboring chromatin. In addition to complementary docking simulation analyses, our data show that retroviral intasome interactions differ in terms of nucleosome interfaces, leading to different capability to engage compacted chromatin.

## RESULTS

### HIV-1 and PFV intasomes show distinct functional nucleosome interfaces *in vitro*

Previous structural studies have shown that the PFV intasome targets the nucleosome by interacting with H2A/H2B dimers and nucleosomal DNA (7). In turn, HIV-1 IN has been presumed to bind other histone components, such as the histone H4 tail (8, 9), suggesting that both intasomes may interact with distinct regions of the nucleosome. To investigate this, we analyzed and compared the structure of the HIV-1 and PFV integration products obtained after integration onto mononucleosomes (MNs) assembled with the commonly used Widom 601 sequence previously shown to provide suitable integration substrates for both models (10).

As shown in **Fig 1A**, both intasome types assembled with their cognate FITC-coupled viral DNA were efficient at integrating into MNs. An analysis of the migration profile of the integration products using polyacrylamide gels clearly revealed distinct insertion profiles for the two intasome types (**Fig 1A**). A comparison of the distance of migration of the integration products with the size ladder (**Fig 1B**) confirmed the difference in the insertion sites for both intasome types. Cloning and sequencing of the integration products (**Fig 1C**) first confirmed the previously reported main PFV integration sites (7), consistent with integration into the exposed major groove of the nucleosomal superhelix location (SHL) ± 3.5 positions, approximately 36 bp away from the dyad. Notably, under our conditions, an additional insertion site could also be observed for this intasome at ∼41 bp and ∼101 bp (**Fig 1C**), which was also observed as a secondary product in analyses of gels from the literature (7). In contrast, for HIV-1, a major insertion site was detected in a different region of the nucleosome, which was found mostly between 51∼56 bp (**Fig 1C**). These insertion sites were mapped to the nucleosome structure reported in **Fig 1D**, confirming that the two intasomes functionally dock and engage distinct regions of the nucleosome to accomplish the integration reaction *in vitro*.

**Figure 1.**
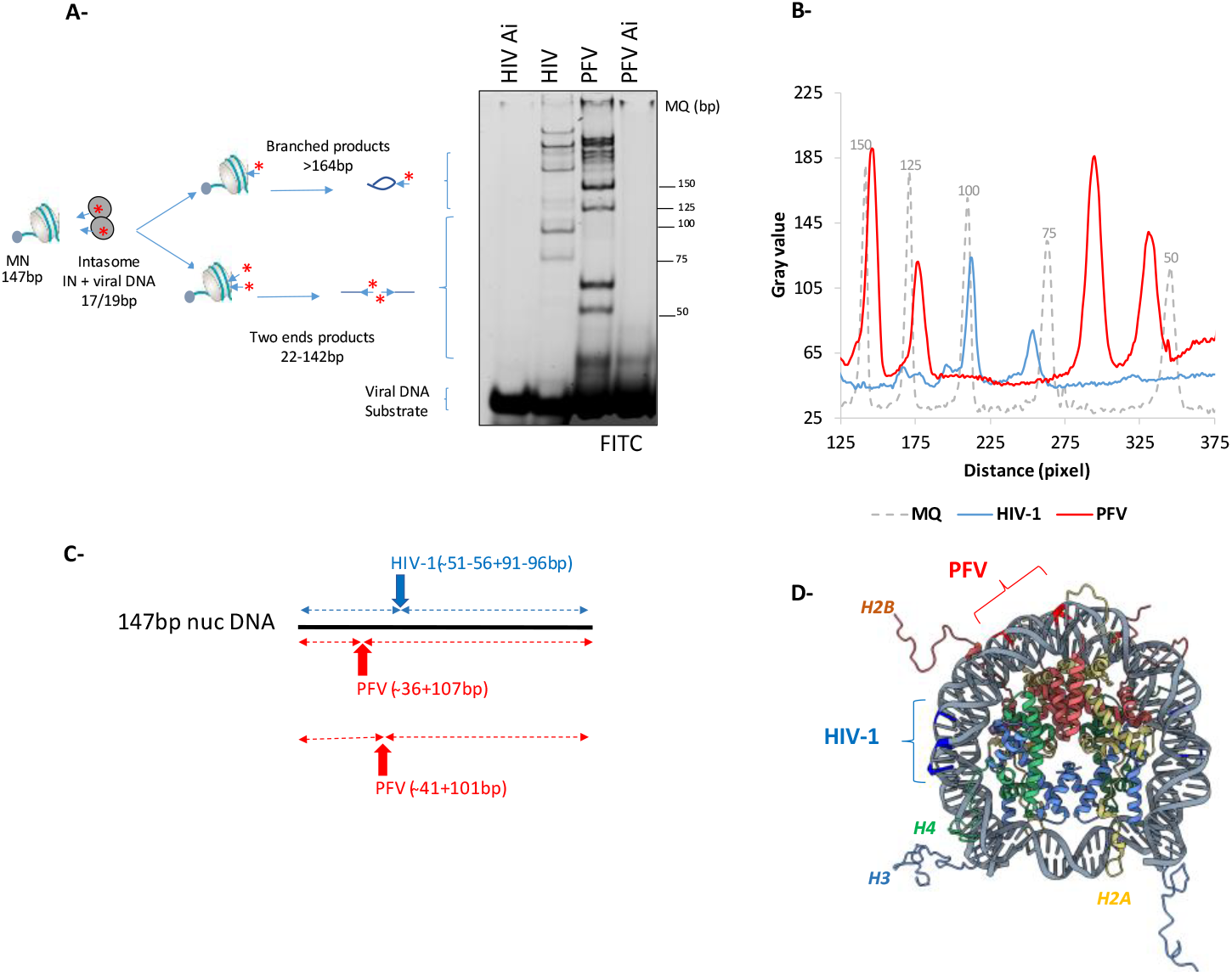
*in vitro* integration selectivity of HIV-1 and PFV intasomes onto nucleosomes. Integration assays were performed using assembled FITC-labelled intasomes and Widom 601 nucleosomes. Integration products were analyzed on 8% polyacrylamide gels, visualized by fluorescence scanning or SYBR safe staining (**A**). Distances of migration of the integration products were compared to molecular weight markers loaded under similar conditions (**B**). Integration products obtained in (**A**) were cloned in bacteria and sequenced. The position of the main HIV-1 and PFV insertion sites were reported onto the 147bp linear Widom 601 sequence (**C**). The corresponding insertion sites were subsequently located on the 3D structure of the 601 nucleosome (**D**).

These differences may be explained by different sensitivities toward nucleosomal structural constraints directed by the interactions between INs and specific histone components and/or DNA structures. In particular, HIV-1 integration relies on the functional association between the intasome and histone tails, which may participate in the stabilization of the TCC and STC as well as their activation (9, 11). To determine whether the selection of the insertion sites at the surface of the nucleosome was dependent on this functional IN/histone association we compared HIV-1 integration on either native or tailless nucleosomes assembled on a typical Widom 601 DNA fragment and recombinant histone octamers. Our results indicate that the deletion of the histone tail significantly decreased the integration efficiency (**Fig 2A-B**), as previously reported (11), probably due to intasome stabilization by histone tails (9). Nevertheless, the integration sites were located at similar positions in both types of nucleosomes (**Fig 2C**).

**Figure 2.**
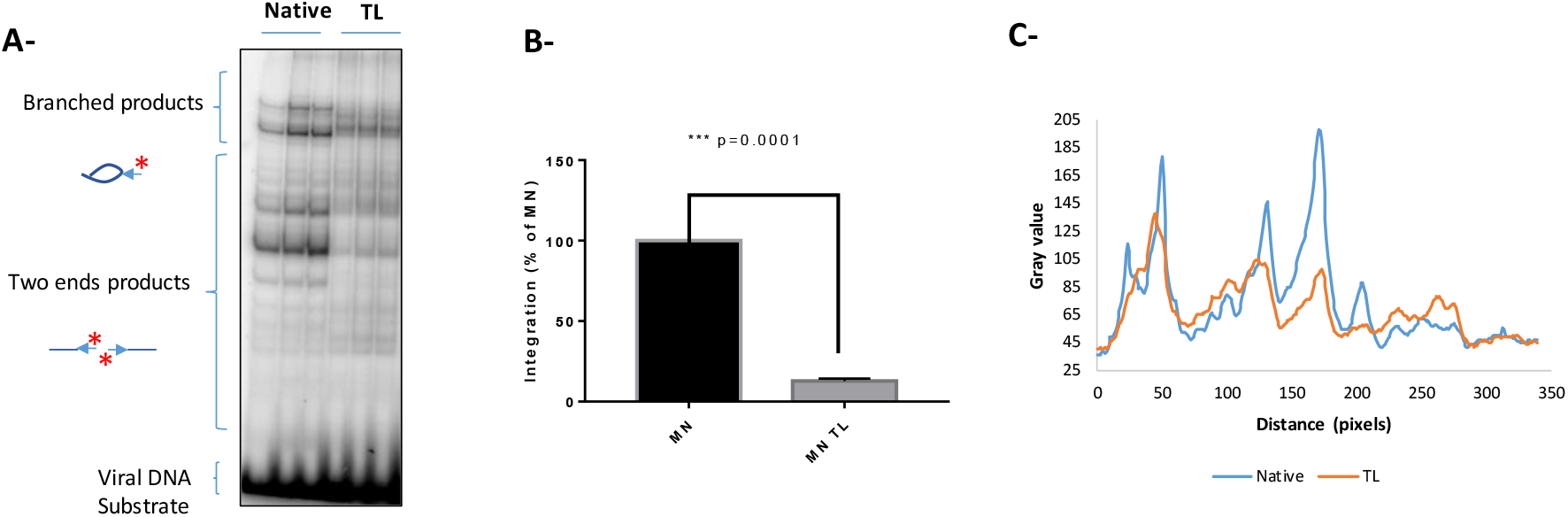
Effect of histone tails on HIV-1 insertion site selection at the surface of the nucleosome. Integration assays were performed using radiolabeled assembled HIV-1 intasome and either native or 601 nucleosomes. Integration products were loaded on 8% native polyacrylamide gels (**A**) and integration was quantified after autoradiography from 3 independent experiments, data are reported as means ±SD (**B**). A t-student statistical analysis was performed to determine the significance of the quantified differences. Insertion site positions along the nucleosome surface was determined by comparison of the distance of migration detected on gel (**C**).

These data indicate that histone tails are not required for the selection of the nucleosomal integration site but more probably important to stabilize the functional incoming STC complex for integration as supported by the previously reported stimulatory effect of the H4 tail (9). Our data also indicate that the difference in nucleosome anchoring between PFV and HIV-1 is probably not due to differential interactions with histone tail components of the nucleosome. In turn, our results indicate that the binding of the intasome to nucleosomal DNA is a major determinant of integration site selection. Structural differences between retroviral intasomes, especially their quaternary structures, as previously reported (6), may also explain the differences in nucleosomal anchoring caused by differences in steric hindrance. To investigate these points, we further compared the physico-chemical properties of nucleoprotein complexes observed after integration onto nucleosomes.

### *In vitro* integration of HIV-1 occurs in highly ordered insoluble nucleocomplexes that are not part of aggregates

To better characterize the nucleoprotein complexes involved in retroviral integration into nucleosomes, we first performed solubility analysis of the integrated products. For this purpose, we performed integration assays followed by sequential centrifugation, and quantification of the soluble fractions. As reported in **Fig 3A**, centrifugation of the HIV-1 samples after integration followed by deproteinization of the supernatant led to the total loss of the integration products. The solubility of the STC-nucleosome complexes (STC-nuc) containing the integration product could be recovered after the addition of 150 mM NaCl, leading to the recovery of most of the integration products, as observed on the polyacrylamide gel (**Fig 3B**). In contrast, higher amounts of PFV STC-nuc were soluble under all the conditions after centrifugation, highlighting their higher solubility under the reaction conditions (**Fig. 3C**, see all the quantifications in **Fig 3. D**).

**Figure 3.**
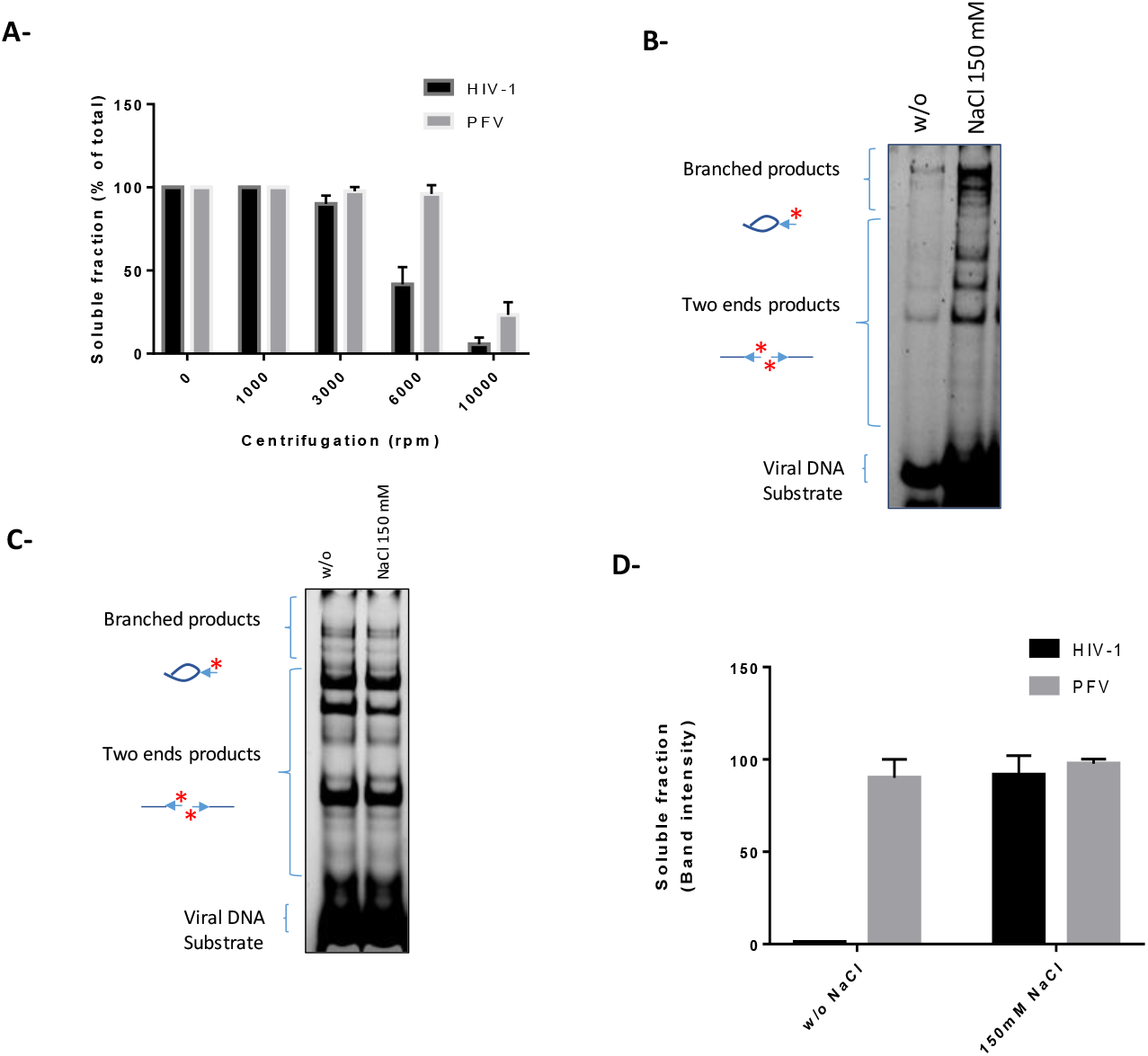
Solubility of the HIV-1 and PFV STC *in vitro*. Concerted integration onto 601 MN was carried with either radiolabeled HIV-1 or PFV intasomes and integration products were centrifugated at the reported speed. Soluble fractions were then deproteinized and the radioactivity was quantified with scintillator counter in both pellet and supernatant (**A**). Results are reported as the % of radioactivity measured in the supernatant versus the total of radiolabeled substrate. Data are reported as means of 3-4 independent experiments ±SD. Concerted integration was then performed similarly with addition of 150mM NaCl into the reaction products before centrifugation at 6.000 rpm. Supernatants from HIV-1 (**B**) and PFV (**C**) were then deproteinized, precipitated and loaded on 8% polyacrylamide gels and autoradiographed (quantification in **D**). Data are reported as means from at least 3 independent experiments ±SD

These data suggest that the loss of HIV-1 integration products during centrifugation performed after integration was mostly due to the insolubility of the nucleosomal functional STC-nuc complexes in contrast to PFV STC-nuc, which remained soluble. The high-order HIV-1 non soluble complexes may be driven by previously reported intasomes stacks (12). Indeed, primate lentiviral intasomes, including HIV-1, ten to fo “p oto-int so e st cks” of ying engths comprising the octameric repeating unit (13, 14). These stacks are formed by domain swapping, which can be limited by the addition of an excess of C-terminal domain (CTD) of IN (15). To determine whether the high-order complexes carrying the nucleosomal integration products could be due to these intasome-mediated stacks, we tested the effect of the addition of purified CTD on nucleosomal activity. As reported in **SI1**, the addition of the HIV-1 CTD did not decrease the integration efficiency but, in contrast, promoted the reaction. This stimulation was not due to an unspecific effect of the CTD since the addition of this domain to the PFV integration reaction led, in contrast, to the inhibition of the efficiency of viral insertion into the nucleosome (**SI2**), probably due to the CTD DNA binding property, which may lead to its binding to the nucleosomal DNA instead of binding to the unrelated PFV intasome.

Taken together, these data indicate that, following the reaction, HIV-1 integration occurs *in vitro* in large functional nucleoprotein complexes, embedding the integration products as highly ordered structures that are not bridged by intasome stacking. As these large complexes may involve multiple intasome-nucleosome interactions, we next investigated the possibility of a multivalent nucleosomal coordination process using nucleosome bridging assays.

### HIV-1 STC bridges several nucleosomes *in vitro* under catalytic conditions

To better understand how nucleosomes are functionally coordinated within retroviral STC, we set up the AlphaLISA assay reported in **Fig 4A**, using nucleosomes coupled to donor and acceptor beads and intasomes. As reported in **Fig 4B**, while no interaction signal between beads was detected without the intasome, the addition of the HIV-1 intasome under reaction conditions induced a strong and reproducible interaction. In contrast, the PFV intasome did not induce any significant interaction signal between beads. Furthermore, experiment performed with IN protein only instead of intasomes did not show an nucleosomal interaction signal, indicating that the binding between MNs is not induced by nonspecific aggregation due to incorrectly assembled components of the intasome (**Fig 4C**). The HIV-1 IN LEDGF/p75 cofactor used in the intasome assembly was also able to induce a bridging signal alone, suggesting that nucleosome coordination may also be mediated by this factor. Since previously observed intasomes stacking may also lead to bridging signals, we tested the effect of adding increasing amounts of CTD to limit these *in vitro* nonphysiologically formed stacks. As reported in **Fig 4D**, the addition of purified CTD did not affect the bridging signal, confirming that the interaction observed is mostly due to the coordination of multiple nucleosomes to a single intasome or STC rather than several clusters of nucleosomes from multiple bonds to multiple intasomes stacked together.

**Figure 4.**
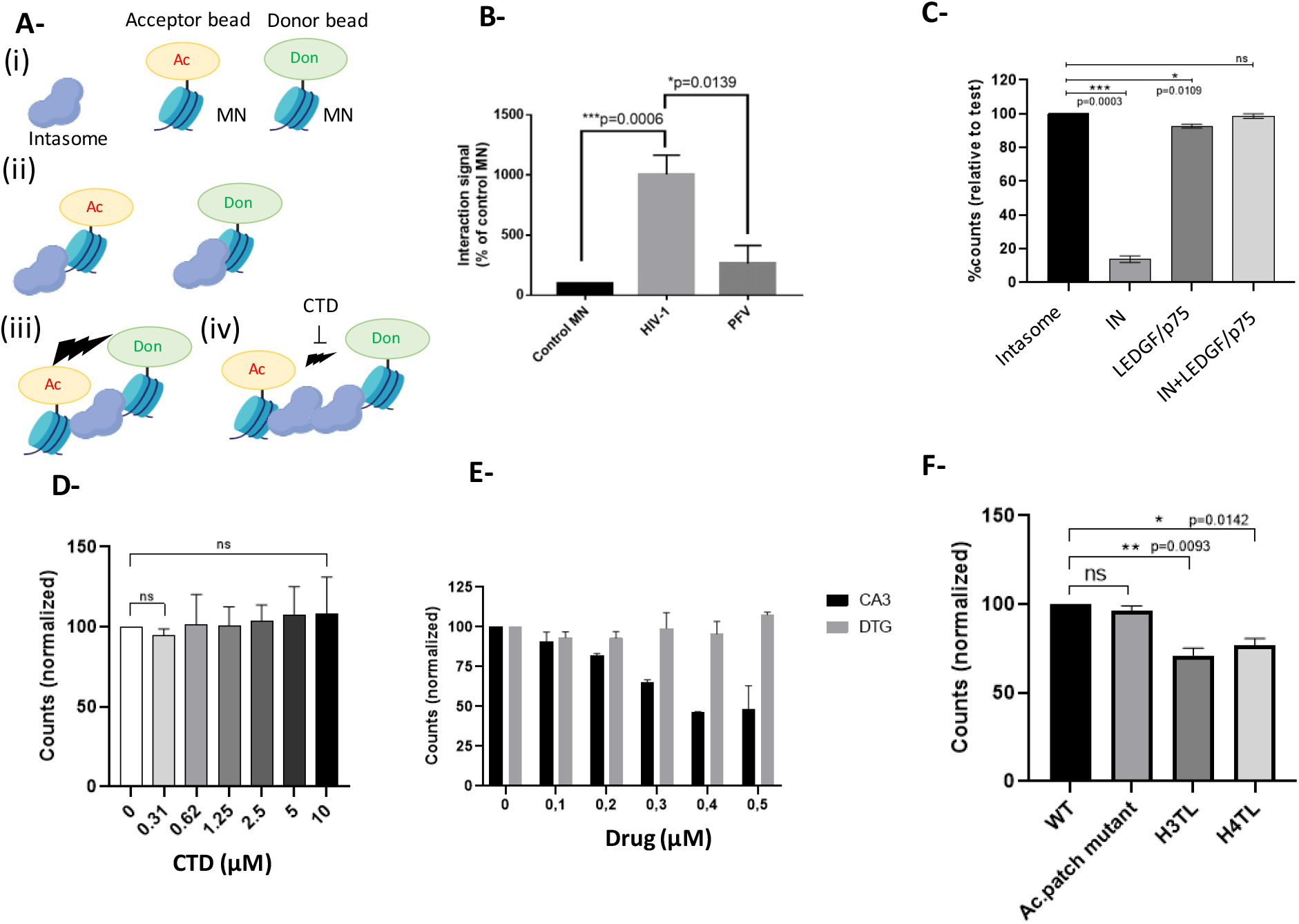
*In vitro n*ucleosome bridging by retroviral intasomes. The AlphaLisa-based bridging assay is schematized in (**A**). Retroviral intasomes are incubated with a mixture of MN coupled to donor and acceptor beads (i). If only one nucleosome is coordinated within on intasome no AlphaLisa signal would be detected (ii). In contrast, the bridging of at least to nucleosomes within either a stack, or an aggregate of intasome, as well as within one intasome forming a TCC will lead to a detectable signal (iii). Addition of HIV-1 CTD can affect the bridging signal mediated by intasome/intasome stacking interactions (iv). Bridging assay has been carried without or in the presence of HIV-1 or PFV intasome under optimized setup conditions (**B**). The bridging assay has been performed with the isolated HIV-1 intasome components with similar concentrations when intasome was used (**C**). Increasing concentration of 220-270 HIV-1 CTD (**D**) or CA3 and dolutegravir drugs (**E**) were added in the MN/HIV-1 intasome bridging assay. Nucleosomes carrying either an acidic patch H2AE92K mutation or deletion in N-terminal histone tails were used (**F**). Data are reported as the means of at least 3 independent experiments ±SD. T-student tests have been performed.

To further investigate whether nucleosome coordination is related to integration activity or intasome binding to multiple nucleosomes, we performed a nucleosome bridging assay under noncatalytic conditions using the dolutegravir (DTG) integration inhibitor. As reported in **Fig 4E**, a similar coordination signal was observed in the presence of DTG, indicating that nucleosome coordination was not dependent on integration catalysis by STC but was likely dependent on the intasome binding to the nucleosome and the formation of the TCC engaging several MNs.

Previous data have shown that binding of the intasome to the nucleosome may involve interactions among IN, LEDGF/p75 and histone tails (8, 9). These histone tails may also be important for this multivalent coordination. To investigate this, we first used CA3 compounds previously shown to bind histones (16) and reported to inhibit the functional association of the intasome with the nucleosome (17). In contrast to DTG treatment, CA3 inhibited the bridging signal, supporting the importance of histone tails in the bridging process (**Fig 4E**). Deletion of H3 or H4 also induced a decrease in the bridging signal confirming that these histone regions are involved in optimal nucleosome bridging induced by HIV-1 intasome binding (**Fig 4F**). Furthermore, additional stacking between adjacent nucleosomes bridged by the intasome may accentuate the interaction signal. Such stacking is known to involve H4 tail interactions with acidic patches (18). We thus further investigated whether this acidic patch nucleosome stacking may be responsible for the detected bridging. We used the H2A E92K acidic patch mutant known to prevent H4 binding and nucleosome interactions (19). As reported in **Fig 4F** the bridging signal was unaffected by the H2A E92K mutation.

Taken together, these data suggest that a significant proportion of HIV-1 TCCs can bridge at least two nucleosomes under reaction conditions. These results also raised the question of the influence of adjacent nucleosomes on the integration reaction.

### HIV-1 and PFV intasomes are distinctively affected by neighboring nucleosomes

Contact with adjacent nucleosomes may affect access to the target catalytic nucleosome and, thus, integration sensitivity toward chromatin compaction. In particular, the distinct nucleosome interfaces found between PFV and HIV-1 intasomes may account for their previously observed differences in sensitivity toward chromatin structure, as demonstrated by integration into reconstituted polynucleosome templates (6, 8, 20).

A comparison of *in vitro* integration of HIV-1 and PFV into isolated nucleosomes and chromatin fibers reconstituted *in vitro* in the presence of the H1 histone linker (see **Fig 5A** for a description of the substrate) revealed that HIV-1 integration is strongly prevented in compacted chromatin. In contrast, the chromatin fiber structure was more efficient for *in vitro* PFV integration than the MN structure was (**Fig 5B**). These data further confirmed the distinct chromatin compaction preferences of the two types of intasome.

**Figure 5.**
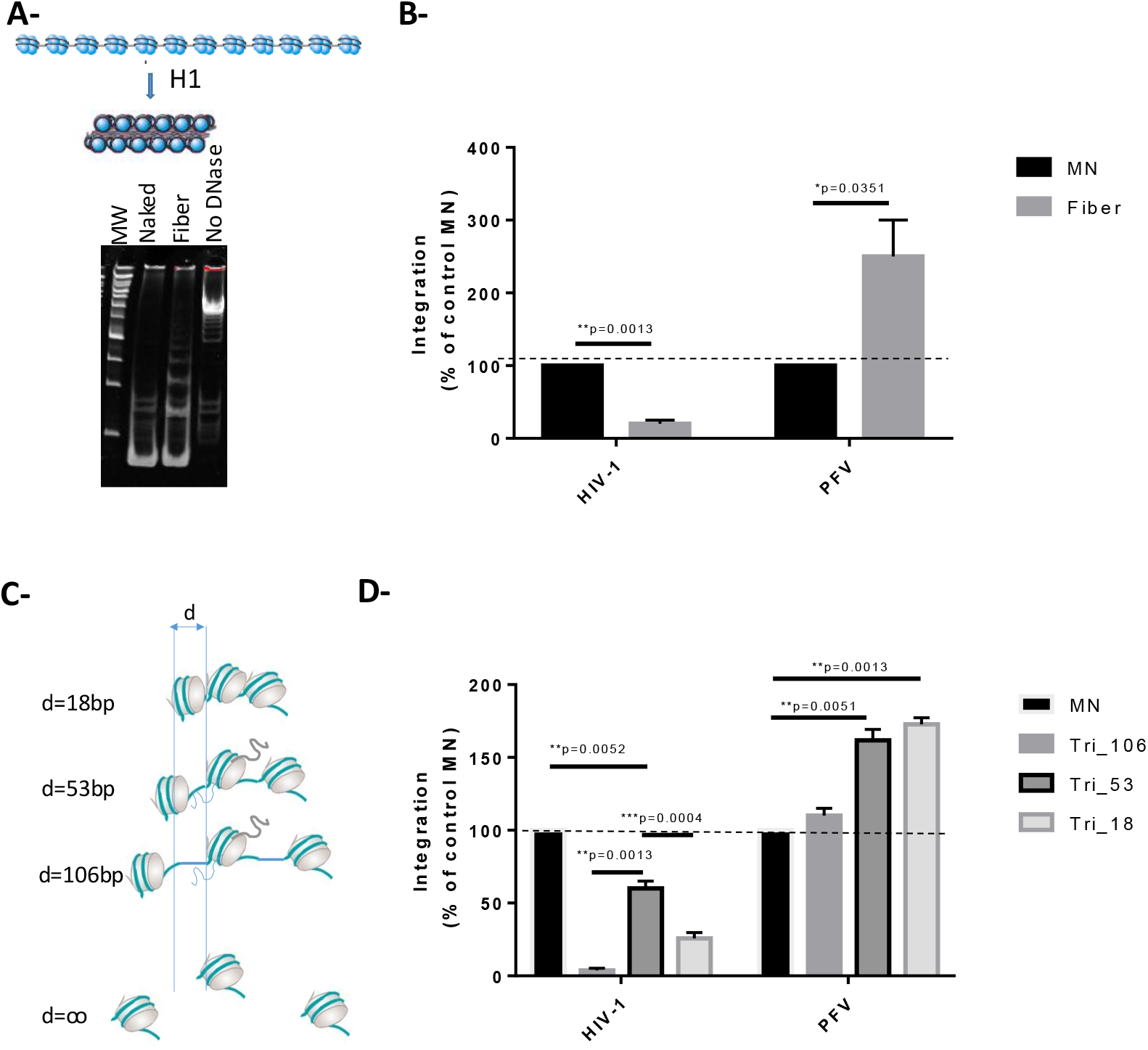
Impact of adjacent chromatin on HIV-1 and PFV integration. Integration was performed in chromatin fiber containing an array of 12 601 sequences assembled in the presence of histone H1. Assembly was monitored by MNase protection after 2 minutes treatment of assembled fiber and corresponding naked DNA (**A**). The efficiency of HIV-1 and PFV *in vitro* integration into naked DNA or fiber was compared by beads integration assays, data are reported as percentage of integration into chromatin fiber, integration into MN being setup at 100% ±SD (3 independent experiments) (**B**). T-student test have been performed to evaluate the significance of the differences between conditions. MN or the trinucleosomes reported in (**C**) were then compared by beads integration assay using either PFV or HIV-1 intasomes. Data are reported as the means percentage of integration in the tri nucleosomes structures, integration into MN being setup at 100% ±SD (3 independent experiments). Data are reported as the means percentage of integration in the chromatin fiber, integration into MN being setup at 100% ±SD (3 independent experiments). T-student statistical tests have been performed to compare each conditions.

The availability of functional intasome/nucleosome interfaces can be modulated differently by the compaction of chromatin. To better investigate this hypothesis, and to more accurately study the impact of neighboring chromatin, we next compared integration onto MN and trinucleosomes (Tri-nuc) structures with distinct internucleosome linker sizes, allowing us to directly monitor the regulatory role of neighboring nucleosomes and chromatin compaction, as illustrated in **Fig 5C**. As reported in **Fig 5D**, quantification of the total amount of HIV-1 integration catalyzed into either the isolated MNs or Tri-nuc showed that the HIV-1 integration efficiency of Tri-nuc was lower than that of MNs. This confirmed that HIV-1 integration is globally inhibited by chromatin compaction.

However, inhibition by compaction was not linear since the integration efficiency of Tri-nuc containing a 53 bp DNA linker (Tri_53) was significantly greater than that of a Tri-nuc with longer (106 bp) or lower (18 bp) linker sizes. This finding suggests that, within compacted structures, specific compaction degrees may be favorable for HIV-1 integration highlighting the role of neighboring nucleosomes in modulating the formation of the functional nucleosomal STC. In contrast, PFV integration was found to accommodate compacted chromatin more efficiently than HIV-1, and was even stimulated on more compacted structures (**Fig 5B**). This highlights the difference in chromatin structure preference for each intasome and underlines the differential behaviors of the two intasome types in terms of chromatin structure.

### Neighboring nucleosomes modulate accessibility to functional HIV-1 interfaces

Taken together, these data indicate that both intasome types are affected differently by chromatin compaction in polynucleosome structures and strongly suggest that neighboring nucleosomes modulate access to functional nucleosome interfaces for efficient integration. We next further investigated the molecular mechanism responsible for these differences. For this purpose, we analyzed the structure of the integration products on native acrylamide gels and quantified the proportion of integration into either the apical adjacent nucleosome or the central nucleosome.

The trinucleosome with 53 bp linker was found to be the best polynucleosomal substrate for HIV-1 integration in previous assays and was, thus, used to determine the insertion site preference in compacted structures. As reported in **Fig 6A**, analysis of the structure of the integration products on polyacrylamide gel revealed only two main full-site integration products, indicating that the HIV-1 intasome catalyzed the integration into one major insertion site in the central region of the trinucleosome (see the migration distance analysis in the right panel). The integration sites were found to shift from approximatively the size of the 601 nucleosome assembly sequence, confirming the insertion into the central region of the acceptor DNA. In contrast, PFV integration into trinucleosome structures led to a similar integration site as that found in MNs but with a 53 bp shift corresponding to the linker size (**Fig 6B**). Positioning of the insertion sites on the trinucleosomal fragment confirmed that the integration occurred mostly on each nucleosome of Tri-Nuc, leading to multiple insertion bands located in the mononucleosomal regions. To confirm these data, we performed an integration assay on beads followed by *EcoR*I cleavage, as reported in **Fig 6C**. The data showed that while most of the HIV-1 integration signal remaining in the bead fraction strongly decreased, PFV integration was less affected, leading to a 70% decrease in the signal. This result is consistent with integration occurring with similar efficiency on each nucleosome for PFV in contrast to HIV-1 (**Fig 6D**).

**Figure 6.**
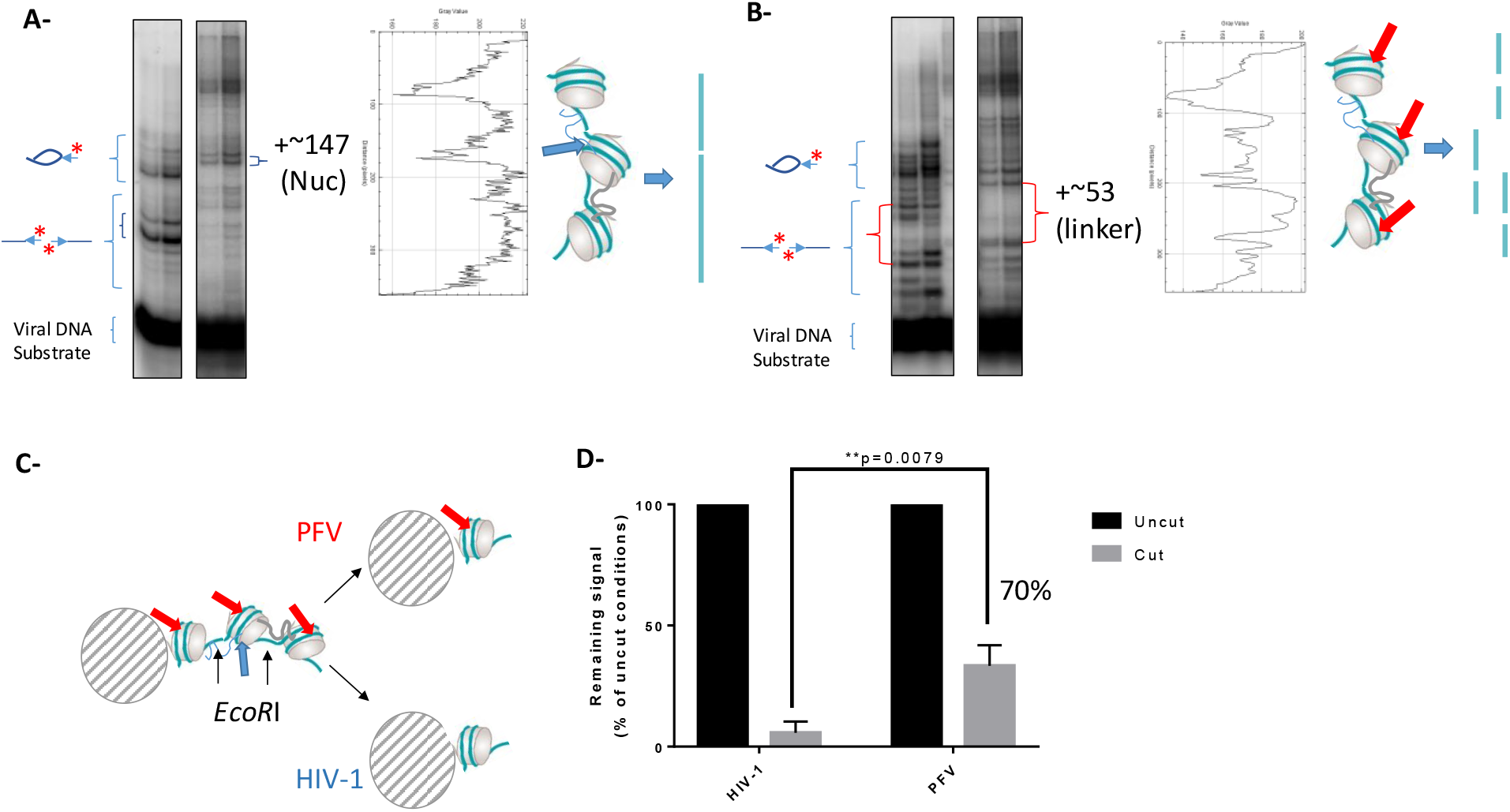
HIV-1 and PFV integration site in mono or trinucleosome. Integration was carried using either the isolated MN or the trinucleosome with 53bp linker and HIV-1 (**A**) or PFV (**B**) intasomes before deproteinization and loading into 8% polyacrylamide gel. Positions of the different integration products on MN are reported on the left side of the gels. The position of the integration product on the trinucleosome surface was estimated from evaluation of the distance of migration using ImageJ software (right panels) and the expected integration products are indicated on the right side of the gels. Beads integration assay was performed followed by *EcorR*I cleavage and quantification of the integration remaining on beads (**C**). Data are reported in (**D**) as the means of 3 independent experiments ±SD and t-Student test.

The localization of the HIV-1 integration site in the trinucleosomal structure at the central nucleosome indicates that only one HIV-1 functional interface is available on this specific trinucleosomal structure and located on the central nucleosome, which is, thus, asymmetrically used as catalytic substrate. These findings also indicate that adjacent nucleosomes are not or are poorly used by HIV-1 integration complexes, suggesting that these nucleosomes may serve for the structuration of the final functional integration complex. This is further supported by the fact that a decrease, or increase, in the linker size between the nucleosomes led to a decrease in the efficiency of integration (cf **Fig 5**). Taken together, these data suggest that while the central nucleosome may be the preferred substrate for the HIV-1 intasome, additional contacts with neighboring nucleosomes may be required for efficient integration into compacted chromatin and specific polynucleosomal structures in agreement with the bridging property of HIV-1 IN shown in **Fig 4**. The positioning of PFV integration sites is consistent with insertions in all the nucleosomes constituting the trinucleosome, indicating that PFV nucleosomal integration is not inhibited by the compaction of the chromatin or by neighboring nucleosomes. Alternatively, our results indicate that the functional PFV/nucleosome interfaces remain accessible despite chromatin compaction unlike those of HIV-1.

### Molecular docking of retroviral intasomes in complex with trinucleosomes

To better understand the interactions between both PFV and HIV-1 incoming intasome on compacted chromatin structure and investigate further the differences found between these functional interactions, we established a model for the intasomes binding to the Tri-nuc structure. For this purpose, HADDOCK 2.4 was employed to dock both intasomes onto the trinucleosome, specifically at the central single nucleosome and at the adjacent double-nucleosome interaction site. A computational model of the complex was first created. The model for the nucleosome was obtained from the 1KX5 structure (resolution of 1.9 Å) (21), which is available on the Protein Databank. The trinucleosome structure was sourced from the 6L4A PDB entry (12.30 Å resolution) (22), whereas the HIV-1 and PFV intasome models were obtained from the 9C9M (2.01 Å resolution) (15) and 3S3M (2.49 Å resolution) (23) PDB structures, respectively. To compare both types of intasome, we used the catalytic tetramer in both analyses. The representative structures from the best scoring cluster were chosen for this work. This structure was chosen not only because of its position as the best scoring cluster but also because it most closely resembled the nucleosome-intasome interaction, as described by Maskell et al. (24).

**Table 1** reporting the data obtained with PFV intasome shows that the HADDOCK docking results indicate a strong binding affinity at both integration sites, with the adjacent double interaction site having a more favorable score (−44.0 ± 13.8) than the single interaction site (−38.5 ± 0.8).

**Table 1.**
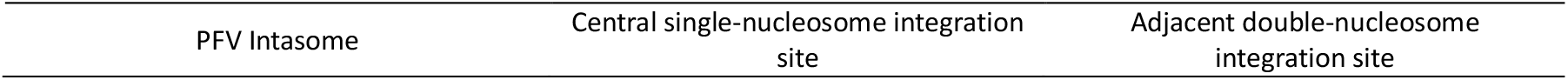
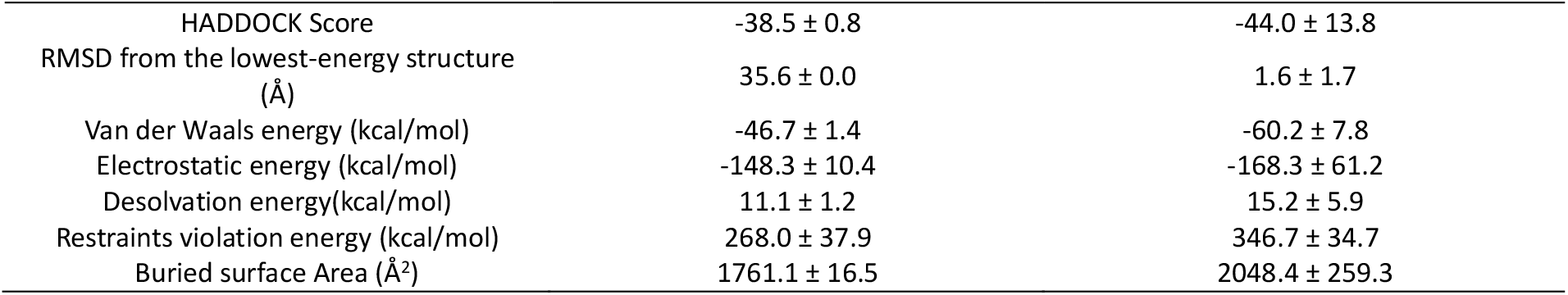
Results of the docking of the PFV intasome with the single and double-nucleosome integration sites of the human trinucleosome.

The lower standard deviation in the single interaction site suggests that the binding poses were more consistent across docking solutions, whereas the higher variability at the double site implies greater structural flexibility. This finding suggests that interactions at the double nucleosome site may provide additional stabilizing contacts, potentially enhancing integration efficiency. With the help of the protein-ligand interaction profiler (25), the interactions between the intasome and nucleosome were analyzed.

In the case of the central single-nucleosome integration site of PFV intrasome, hydrogen bonds are formed between the hydroxyl group of nucleosome H2A S1 and the amine group of the intasome CTD A328, between the guanidinium group of nucleosome H2A R3 and the amide group of the intasome CTD N360, between the guanidinium group of nucleosome H2A R11 and the guanidinium group of intasome CTD R334, and between the hydroxyl group of nucleosome H2B S120 and the amine group of intasome CCD L230. Notably, as reported in the literature, the intasome interacts with the tail of the H2A histone, particularly with R11, which was reported as a key interacting residue by Wilson et al. (26). Furthermore, it also interacts with the C-terminus of the H2B histone, which has also been reported in the literature (24, 26). **Fig 7A** shows the interaction between the PFV intasome and the human nucleosome at the central single-nucleosome integration site. In this figure, the interaction between the H2A histone tail (orange) and the PFV CTD (yellow) is clearly detected. Furthermore, the C-terminal helix of H2B (white) can also interact with the PFV CCD (cyan).

**Figure 7.**
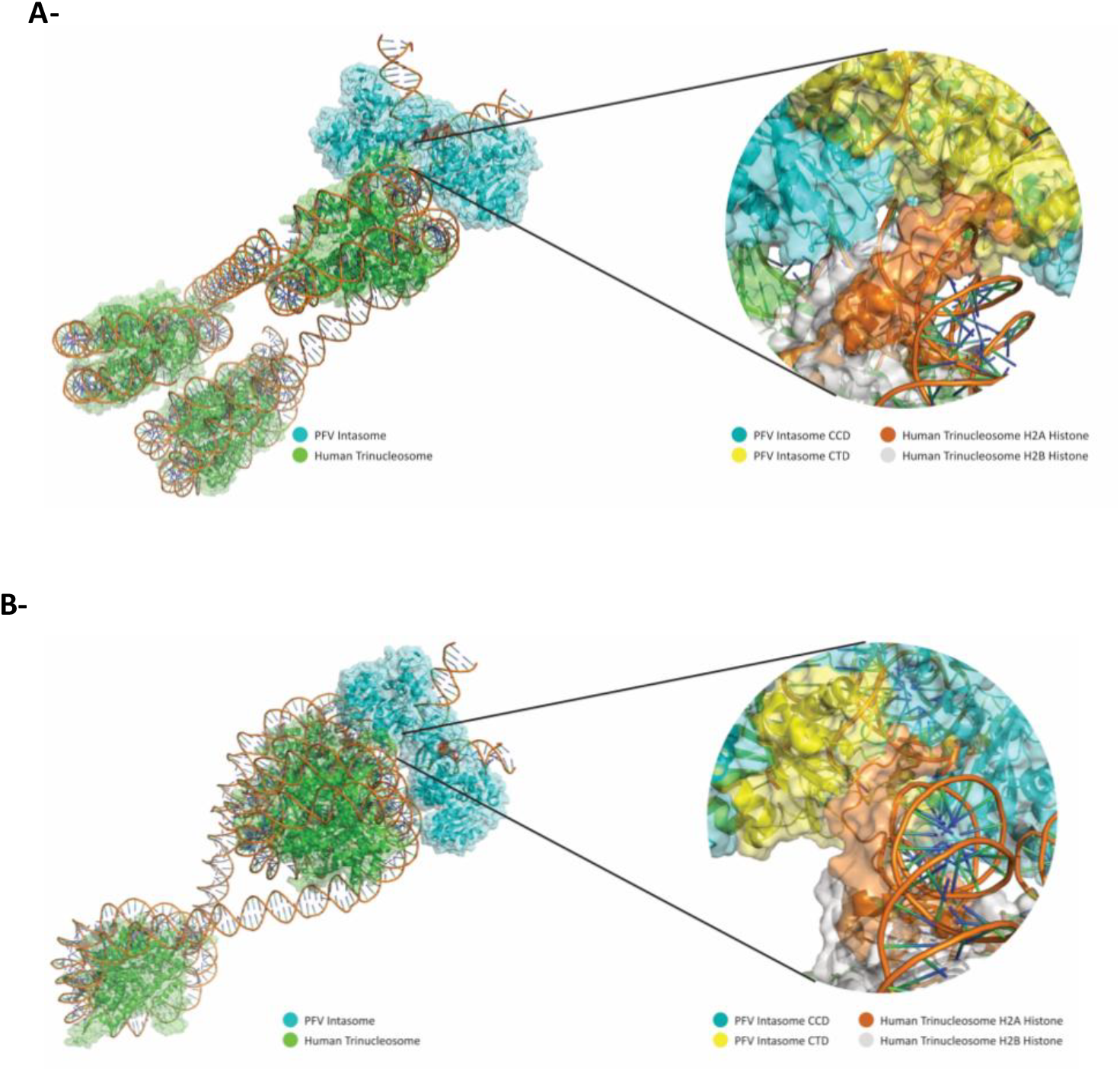
Docking model of the interaction between the incoming PFV intasome and the single-nucleosome integration site (A) or the double-nucleosome insertion site (B) into trinucleosome.

For the double-nucleosome integration site of PFV, hydrogen bonds are formed between the hydroxyl group of the nucleosome H2A S1 and the hydroxyl group of the intasome CTD S332, between the amino group of the nucleosome H2A K9 and the amine group of the intasome adenosine 17, and between the guanidinium group of the nucleosome H2A R11 and the hydroxyl group of the intasome CCD Y212.

**Fig 7B** depicts the interaction between the PFV intasome and the double-nucleosome integration site. Once again, the interaction between the H2A histone tail (orange) and the PFV CTD (yellow) can be predicted despite the proximity of the two nucleosomes. While the H2B C-terminal helix is not shown to be in contact with the CCD, the two domains are still relatively close, not excluding the possibility of interaction between them. Compared with the single-nucleosome integration site, the same area of the nucleosome interacts with the intasome, which can explain why integration is possible at both sites despite the compaction.

The results of the docking of the HIV-1 intasome and the human trinucleosome are displayed in **table 2**.

**Table 2.**
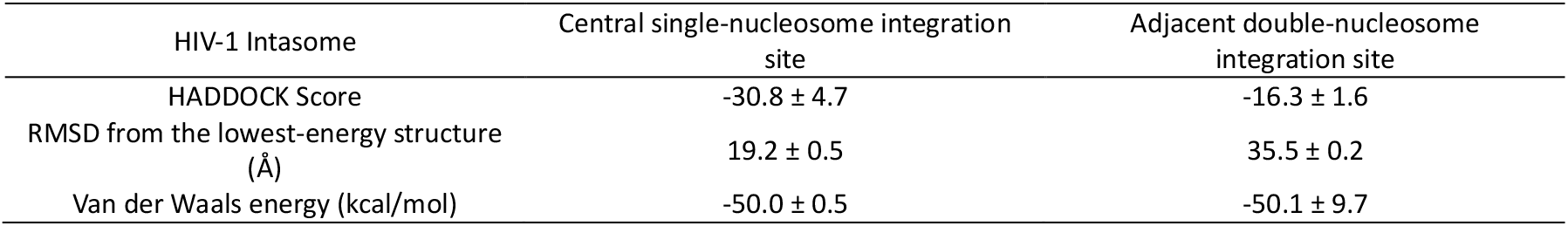
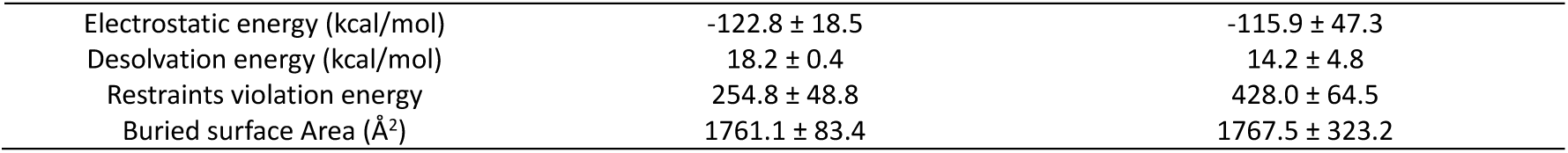
Results of the docking of the HIV-1 intasome with the single and double-nucleosome integration sites of the human trinucleosome.

The single nucleosome complex has a much more favorable HADDOCK score (−30.8 vs. -16.3), indicating that the HIV-1 intasome forms a much stronger and more stable interaction in the single-nucleosome integration site. In the double-nucleosome integration site, the complex has a lower negative score, suggesting that the additional nucleosome might create steric hindrance or make binding less efficient than a single nucleosome interaction. Once again, the interactions were analyzed via protein-ligand interaction profiler (25). These findings strongly support the biochemical results showing that, in contrast to PFV, HIV-1 integration is prevented on a trinucleosomal structure, and the insertion site is re-oriented toward the central nucleosome resulting in a more stable interaction with the incoming intasome.

With respect to the single-nucleosome integration site, hydrogen bonds are formed between the guanidinium group of nucleosome H4 R3 and the amide group of the intasome inner CTD N254, and between the N-terminal amine of nucleosome H4 V21 and the backbone carbonyl group of nucleosome CCD G70. Additionally, salt bridges are formed between the amino group of nucleosome H4 K8 and the carboxyl group of the intasome CCD E170, and between the amino group of nucleosome H4 K16 and the carboxyl group of the intasome CTD D232. These highlight the crucial role of the H4 histone tail and its significant interactions with the intasome CTD (9, 11). Notably, H4 R3 and CTD D232 emerged as key residues in the interaction between the HIV-1 intasome and the human nucleosome (8). **Fig 8A** highlights the association between the incoming HIV-1 intasome and the central nucleosome showing the interaction between the histone H4 tail in red and both the intasome CTD (yellow) and the CCD (cyan).

**Figure 8.**
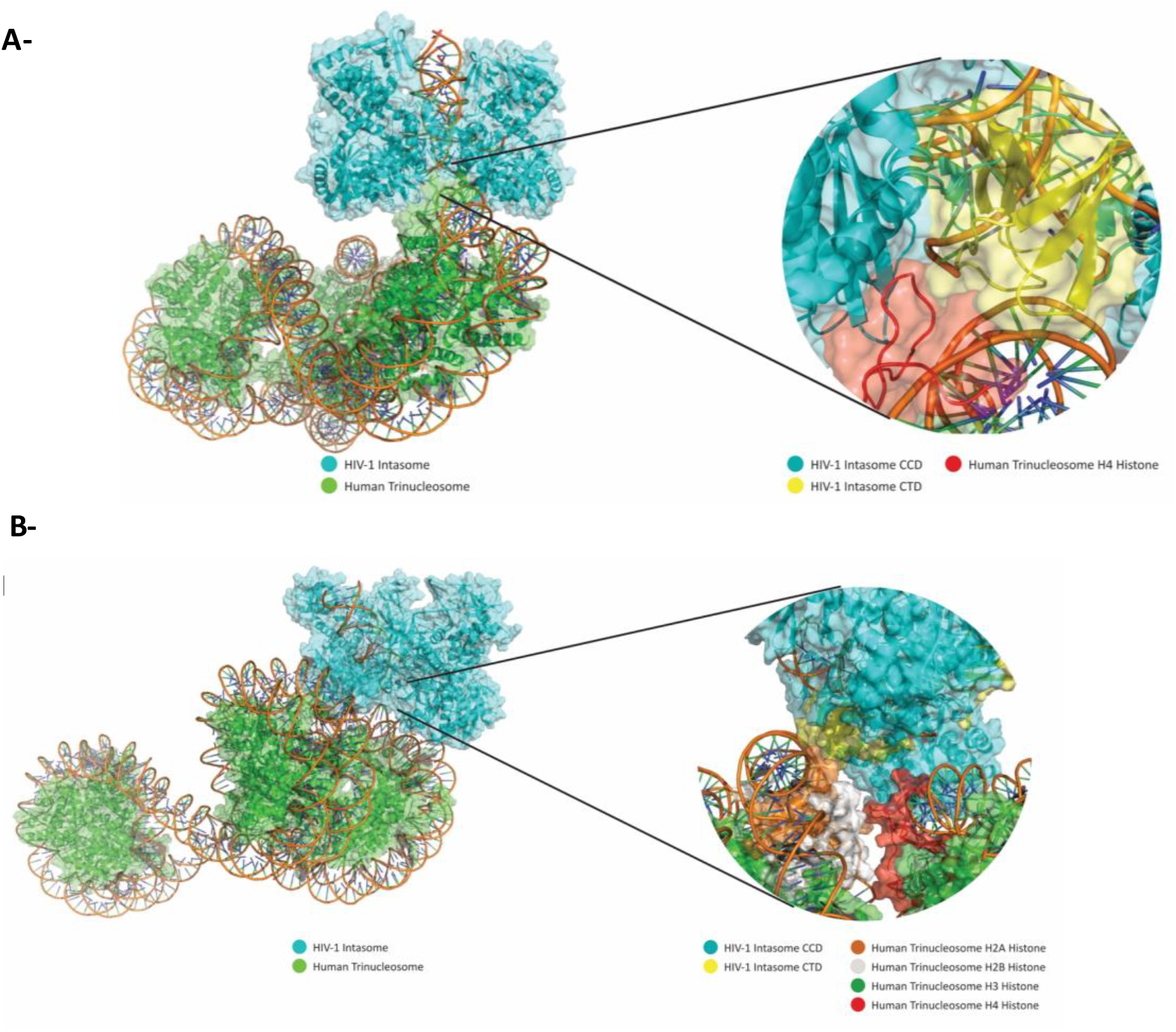
Docking model of the interaction between the incoming HIV-1 intasome and the single-nucleosome integration site (A) or the double-nucleosome insertion site (B) into trinucleosome.

In relation to the adjacent double-nucleosome integration site, only one hydrogen bond was formed between the hydroxyl group of nucleosome H4 S1 and the backbone carbonyl group of the intasome CCD E69 for the target nucleosome. In the adjacent nucleosome at this double-nucleosome integration site, bonds are established between the backbone amine of nucleosome H2A A14 and the carboxyl group of the intasome CTD D256 and between the backbone amine of nucleosome H2A K15 and the backbone carbonyl group of the intasome CTD S255.

**Fig. 8B** shows that the HIV-1 intasome is not able to interact with the adjacent nucleosome in the same manner as it does with the central nucleosome. In addition to the less negative docking score, this result indicates that the intasome is not able to bind to the trinucleosome at the preferred site, interacting more with the H2A histone of the adjacent nucleosome than with the H4 tail of the target nucleosome. In this case, we could expect that the previously reported stabilization and stimulating effects of binding to H4 cannot occur in this situation. This may explain the poor efficiency of integration found for this structure *in vitro*.

## CONCLUSION

Retroviral integration occurs at the surface of a nucleosome in the host chromosome. Nucleosomes are always embedded within different levels of chromatin compaction, prompting the question of whether they engage in long-range interactions or proximal contacts with neighboring chromatin regions. Previous studies have shown that retroviral integration models are differently affected by the degree of chromatin compaction (5, 6). However, the molecular mechanism underlying this difference in sensitivity remains to be determined. This differential retroviral sensitivity toward chromatin structure could be due to the difference in steric hindrance related to the differences found in the quaternary structures of the intasome (4) as well as possible differences in their anchoring interfaces at the surface of the nucleosome, which may be differentially masked by chromatin compaction.

Exploring these hypotheses by comparing two different intasome type, namely PFV and HIV-1, with different chromatin sensitivity patterns both in cells and *in vitro*, revealed that both intasome types bind different regions of the nucleosome, leading to different insertion sites within the corresponding STC. Indeed, while PFV integration occurs mostly on the previously reported IS (7) analysis of the HIV-1 insertion sites revealed that this intasome preferentially binds the 51∼56bp of the nucleosomal DNA. Masking this insertion site in compacted chromatin would lead to a decrease in integration efficiency, as previously observed. Testing this hypothesis by positioning the HIV-1 and PFV nucleosomal insertion sites identified on the tri-nucleosome compacted structure revealed that while the PFV nucleosomal insertion site was accessible on all three nucleosomes, only one HIV-1 IS appeared exposed and suitable for integration and was located on the central nucleosome. In contrast, the corresponding HIV-1 ISs located on adjacent nucleosomes are deeply masked at nucleosome interfaces, preventing integration onto these regions. Molecular docking simulations of intasome docking to these identified sites in the three nucleosomes of a compacted trinucleosome led to interactions models confirming that PFV incoming intasome can efficiently bind all the nucleosomal insertion sites. In contrast, HIV-1 binding to adjacent nucleosomes was clearly unfavored compared with that of the central nucleosome due notably to the masking of both DNA and histone interactions required for efficient integration.

*In vitro* integration assays onto trinucleosomes and chromatin fiber structures support this model, confirming the difference in sensitivity toward chromatin compaction between PFV and HIV-1 and showing that PFV integration accommodates the chromatin compaction more easily than HIV-1 integration does, as suggested by our model. Furthermore, analysis of the trinucleosomal integration products confirmed that the PFV intasome could functionally interact with all three nucleosomes, as expected from our model, and explain its ability to accommodate compaction. In contrast, HIV-1 integration appeared to occur mostly in the central nucleosome region, as expected from the IS masked onto the adjacent nucleosomes, which explains the negative effect of chromatin compaction on this retroviral integration system. These findings provide insights into the molecular mechanisms of retroviral integration. Compared with the HIV-1 intasome, the PFV intasome has a stronger and more stable binding affinity at both central and adjacent nucleosome integration sites. The favorable HADDOCK scores and interaction profiles suggest that PFV intasomes can efficiently integrate at both integration sites. In contrast, the HIV-1 intasome has reduced affinity at the adjacent nucleosomal sites, likely due to steric hindrance surrounding the insertion site and weaker interactions with key histone residues. This may explain why the HIV-1 intasome preferentially integrates at the central nucleosome site and is unable to efficiently target each nucleosome within a compacted trinucleosome.

Our analysis also revealed that the ability of reducing the nucleosome linker size to inhibit integration was not linear. Indeed, we unexpectedly found that certain lengths of nucleosome linkers favored integration within the set of poly nucleosomes used in our assay. However, the integration efficiency detected in these particular polynucleosomes never reached the level found in mononucleosomes, which may recapitulate the most relaxed chromatin models. Bringing closer to or moving away from the neighboring nucleosomes by modifying the nucleosome linker was indeed found to affect the efficiency of integration. These original results may suggest additional contacts between STC components and adjacent nucleosomes while integration would be performed mostly onto the central nucleosome. A nucleosome-bridging Alphalisa assay (NBA, set up in our work) confirmed that, in contrast to PFV, HIV-1 efficiently bridges at least two different nucleosomes. Even if direct contact between nucleosomes cannot be fully ruled out, such nucleosome-nucleosome interactions are not supported by our data obtained with acidic patch mutants. Additionally, this bridging property was probably not due to IN/IN aggregation or intasome/intasome contacts previously shown to induce their oligomerization since the addition of the HIV-1 CTD did not affect the interaction signals. In contrast, omitting the LEDGF/p75 protein largely decreased the association signal as well as the deletion of H3 or H4 tails. The use of LEDGF/p75 alone confirmed the ability of this factor to bridge several nucleosomes. This property may, thus, participates in the global nucleosome bridging ability of the HIV-1 intasome. These results may be reminiscent of the previously reported multivalent recognition of nucleosomes by LEDGF/p75(27), which involves multiple interactions between the PWWP domain of the factor and histone tails and nucleosomal DNA. In particular, in addition to the PWWP interaction with the histone tail, an additional region of interaction was found in the region of the histone octamer acidic patch, and the DNA interface remained dynamic in the context of dinucleosomes. Additionally, specific sites within the intrinsically disordered central region of LEDGF/p75 selectively interact with nucleosomal or extranucleosomal DNA. In concert with our data, these results strongly suggest that the intasome bridging property mostly relies on the LEDGF/p75 cofactor and histone tail interactions, which may be involved in the contact of the HIV-1 intasome with adjacent nucleosomes and the selection of suitable local chromatin structures suitable for integration, as proposed in the model reported in **Fig 9**.

**Figure 9.**
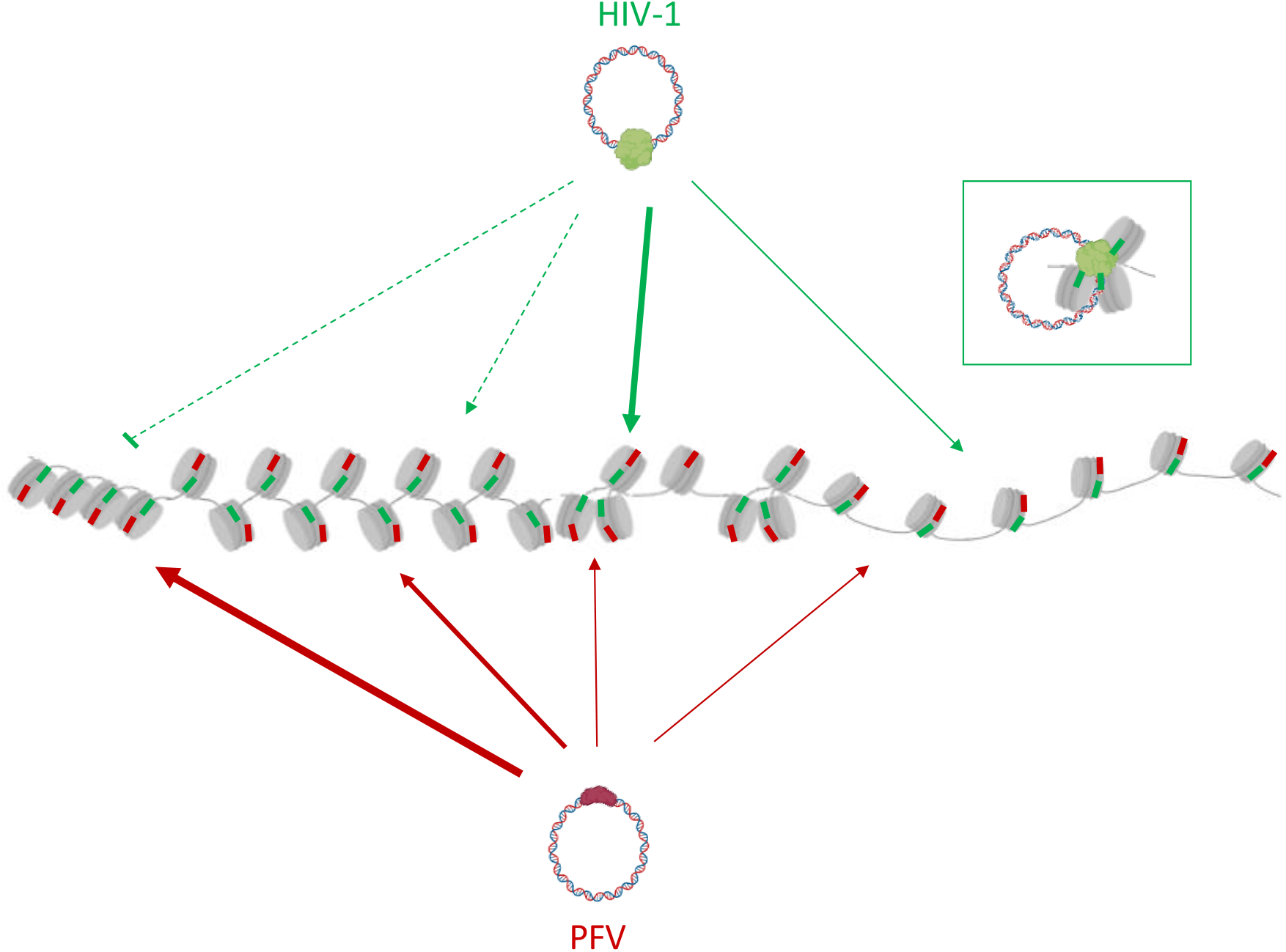
Model for local chromatin structure scanning by HIV-1 incoming intasome. After a global targeting toward specific chromatin region the efficiency of retroviral nucleosome binding would rely on the local availability of the related insertion site govern by the level of chromatin compaction. For PFV the functional interface of association with the nucleosome would be accessible in most of the chromatin structures. Compaction of the chromatin may over-exposed the optimal intasome-nucleosome interface. This would result in a high capability of PFV intasome to accommodate chromatin structures with a preference for compacted nucleosomes. HIV-1 integration would require increased accessibility of the nucleosome interface in open and specifically structured compacted chromatin structure allowing the correct association between the targeted and adjacent nucleosomes (as shown in the focused panel).

In this model, the availability of different retroviral ISs modulates the ability of incoming intasomes to efficiently bind nucleosomes in specific chromatin regions. After global intasome targeting toward specific chromatin regions, multivalent interactions between IN, LEDGF/p75 and both targeted and adjacent nucleosomes allow the selection of suitable local structures with nucleosome density and compaction levels suitable for efficient functional docking of the target capture complex and integration catalysis. In the case of PFV, the increased availability of the nucleosomal IS within chromatin would allow the incoming intasome to accommodate more broad structures and compaction levels. A higher level of compaction may lead to more exposed PFV ISs and fully mask HIV-1 ISs. This mechanism would constitute an additional parameter in the global intasome targeting toward host chromatin during the final local association process.

Our data additionally suggest that the incoming HIV-1 intasome may form a functional STC on multiple nucleosome structures involving at least two structural nucleosomes neighboring the functional one. In this process, LEDGF/p75 may participate in this functional binding in addition to providing a way to gain access to the dynamic transcribed region of the chromatin where the insertion site is available for integration. This multivalent interaction may be important for stabilizing the STC complex, which may be of particular importance in the highly transcribed regions targeted by HIV-1. Previous remodeling activities, such as SWI/SNF or FACT histone chaperones, which have been shown to improve HIV-1 integration (20, 28), may also help in generating functionally suitable local chromatin structures suitable for integration. The recently reported histone chaperone activity of LEDDGF/p75 (Poster communication in CSH retroviruses meeting 2025 from Joshi et al.) may be directly involved in the local remodeling of the nucleosomal insertion site, directly providing the correct local compaction level for integration during the integration process.

The exact role of LEDGF/p75 in this docking process remains to be fully elucidated, and how this factor may influence the structure of the chromatin locally or neighboring it in an integration-coupled manner remains to be answered. Indeed, one limitation of our docking study is the lack of LEDGF/p75 in the structure used, providing the possibility to detect these interactions. Additional protomers of IN are also missing in the docking study, indicating the detection of potential additional contact with adjacent nucleosomes in these analyses, as predicted from our biochemical experiments.

Our model also implies a new notion of the retroviral integration process involving both catalytic and structural nucleosomes. Additional structural approaches, such as cryoelectron microscopy or electron tomography, are thus needed to gain insight into the atomic interaction within this newly identified integration complex and define all the interactions between the intasome and both structural and catalytic nucleosome components. Such structural insights remain largely limited by the heterogeneity of the HIV-1 STC-Nuc preparations, probably due to the multivalent intasome/nucleosome interactions demonstrated in our work. The data reported here will thus help in defining the best polynucleosomal substrate for future attempts to resolve the structure.

Although additional structural data are needed to fully elucidate the molecular determinants of retroviral intasome interactions with chromatin, our findings reveal that different retroviral species have evolved distinct strategies to engage nucleosomes within specific chromatin regions. These preferred nucleosomal targets vary between viruses, reflecting divergent mechanisms to access suitable regions of host chromatin and optimize host genome invasion.

## MATERIALS and METHODS

### Protein purifications and intasome assembly

Integrase of PFV has been purified and the intasome assembled as described in (17). Briefly, PFV IN and its cognate vDNA were mixed and assembled intasomes were purified by size exclusion chromatography. Concordantly with published works, the PFV intasome, which is composed of a tetramer of IN, eluted around 11 mL. The HIV-1 IN was expressed in *E. coli* (Rosetta) and the cells were lysed in buffer containing 50 mM Hepes pH 7.5, 5 mM EDTA, 1 mM DTT, 1 mM PMSF. The lysate was centrifuged and IN extracted from the pellet in buffer containing 1 M NaCl, 50 mM Hepes pH 7.5, 1 mM EDTA, 1 mM DTT, 7 mM CHAPS. The protein was then purified on butyl column equilibrated with 50 mM Hepes pH 7.5, 200 mM NaCl, 1 M ammonium sulfate, 100 mM EDTA, 1 mM DTT, 7 mM CHAPS, 10 % glycerol. The protein was further purified on heparin column equilibrated with 50 mM Hepes pH 7.5, 200 mM NaCl, 100 mM EDTA, 1 mM DTT, 7 mM CHAPS, 10 % glycerol. LEDGF/p75 was expressed in PC2 bacteria and the cells were lysed in lysis buffer containing 20 mM Tris-HCl pH 7.5, 1 M NaCl, 1 mM PMSF added lysozyme and protease inhibitors. The protein was purified by nickel-affinity chromatography and the His-tag was removed with 3C protease, 4°C over night. After dilution down to 150 mM NaCl, the protein was further purified on SP column equilibrated with 25 mM Tris pH 7.5, 150 mM NaCl (gradient from150 mM to 1 M NaCl), then DTT was added to 2 mM final and the protein was concentrated for Gel filtration. Gel filtration was performed on a superdex 200 column (GE Healthcare) equilibrated with 25 mM Tris-HCl pH 7.5, 500 mM NaCl. Two mM DTT were added to eluted protein that was then concentrated to about 10 mg/ml. The C-terminal domain (CTD, residues 220– 270) of HIV-1 Integrase was purified as previously described (Kanja et al., J Virol. 2020 Sep 29;94(20):e01035-20. doi: 10.1128/JVI.01035-20. PMID: 32727879; PMCID: PMC7527040). The CTD was expressed in BL21(DE3) Escherichia coli cells grown in Luria-Bertani (LB) medium supplemented with 50 µg/mL ampicillin and 10% (w/v) sucrose at 37°C with constant shaking at 220 rpm. When the c t e e che n O of .8, exp ession s in ce y ing. isop opy β-D-1-thiogalactopyranoside (IPTG), and incubation continued overnight at 25°C with shaking at 190 rpm. Cells were harvested by centrifugation at 4000g for 20 min at 4°C, resuspended in 10 g/L NaCl, washed, n sto e t − ° efo e p ific tion. ypic y, –6 g of wet cell mass was obtained per liter of culture. For purification, the cell pellet was thawed and resuspended in lysis buffer (25 mM HEPES, pH 8.0, 1 M NaCl, 10 mM imidazole, pH 8.0) at a ratio of 10 mL buffer per gram of cells, with the addition of protease inhibitor cocktail (1 tablet per 50 mL). The cells were lysed by ultrasonication using a 13 mm probe, applying 2-second pulses (on/off) at 40% amplitude on ice to prevent overheating. Cell debris was removed by ultracentrifugation at 185000g for 1 h at 4°C using a Ti45 rotor. The clarified lysate was applied to a 5 mL HisTrap FF Crude column (Cytiva) equilibrated with lysis buffer. Unspecifically bound proteins were removed by a stepwise increase of imidazole concentration: 8 column volumes (CV) at 10 mM imidazole, 8 CV at 20 mM, and 8 CV at 40 mM. The protein of interest was then eluted using a linear gradient from 40 to 500 mM imidazole over 12 CV. Fractions containing the purified protein were pooled and concentrated to 0.5 mL using a 15 mL Amicon Ultra-3K centrifugal filter unit (Millipore) before loading onto a Superdex 75 16/60 gel filtration column (Cytiva) pre-equilibrated with storage buffer (25 mM HEPES, pH 8.0, 1 M NaCl). The appropriate fractions were collected and concentrated to ∼5 mg/mL using a 4 mL Amicon Ultra-3K filter unit, aliquoted and flash-frozen in liquid nitrogen for storage at −8 °.

### Nucleosome assembly

Mononucleosomes were assembled as previously described for chromatin assembly (17). Briefly, 5 µg of biotinylated 147 bp Widom fragment (Epicypher) were mixed with an excess of 10 µg of human native recombinant octamers produced in *E. coli* p ch se f o the “istone o ce” Protein Expression and Purification (PEP) facility from the Colorado State University, https://histonesource-colostate.nbsstore.net) in Tris-HCl pH 7.7 and 2 M NaCl in 100 µl final volume. Salt dialysis was then performed to decrease the salt concentration to 0 using slide-A-Lyzer MINI dialysis device, 7 k MWCO (Fisher Scientific). Assembly was checked by electro-mobile shift assay (EMSA) on 8 % native PAGE stained with SYBR safe and SDS-PAGE stained with instant blue. The acidic patch mutant H2A E92K was purchased at Epicypher (ref. SKU:16-1030).

### Drugs

The CA3 compound was synthetized as previously described (29). Dolutegravir was purchased at INTERCHIM SA

### Integration assays

Integration assays using purified assembled intasomes were performed as previously (17). Briefly, 100 ng of DNA assembled on 100 ng of histone octamers (24 nM final concentration) were incubated with 30 nM final concentration of purified intasome in 100 mM NaCl, 20 mM BTP pH 7, 12.5 mM MgSO_4_ (for PFV intasome), or in 20 mM HEPES pH 7, 7.5 % DMSO, 8 % PEG, 10 mM MgCl_2_, 20 µM ZnCl_2_, 100 mM NaCl, 5 mM DTT final concentration (for HIV-1 intasome) in a final volume of 40 µL. The mix was then incubated at 37°C for 15 mins. Then reaction was stopped by the addition of 5.5 µL of a mix containing 5 % SDS and 0.25 M EDTA and deproteinized with proteinase K (Promega) for 1 hour at 37°C. Nucleic acids were then precipitated with 150 μ of eth no o e night t − °. p es e e then spun at top speed for 1 hour at 4°C, and the pellets were dried and then re-suspended with DNA loading buffer. Integration products were separated on an 8 % native polyacrylamide gel.

For site sequencing, after integration reaction, deproteinization and DNA precipitation PCR was performed using the HU5pro (‘ GTGGAAAATCTCTAGCA ‘ and 601rev ‘ GG G GG G ‘ primers. The PCR products were then purified using the Wizard®SV gel and PCR clean up system (Promega, Ref. A9281) and cloned into pGEM®-T easy vector (Promega, Ref. A1360). Cloned fragments were then sequenced using T7 primer.

### Nucleosome Bridging Assay (NBA)

The NBA was performed on the basis of typical intasome/nucleosome alphaLISA previously setup (17). MN-biotin and MN-DIG (2 nM final concentration) were incubated in buffer NBA (8.1mM Hepes pH7.0, NaCl 100mM, Potassium acetate 13mM, CaCl2 975µM, ZnCl2 3.25µM, DTT 3.25mM, 1.25% DMSO). Streptavidin coated alpha donor beads (Revvity) were added at a final concentration of 5µg/ml, and then incubated at RT with MNs for 5 min before adding the the anti-DIG conjugated AlphaLisa acceptor Beads at the same final concentration in the dark. Short spin was performed at 300rpm and incubation was extended to 10 min at RT in dark. Then 1µl of freshly assembled intasome was added at 16 µM final concentration of Integrase. The plate was incubated at 37°C in a Victor Nivo reader for 15 min, before measurement of interaction signals.

### Molecular docking

To explore how the HIV-1 and PFV intasomes interact with the human trinucleosome, computational models of these complexes were constructed. The trinucleosome structure was sourced from the 6L4A PDB entry (12.30 Å resolution) (22), while the HIV-1 and PFV intasome models were obtained from the 9C9M (2.01 Å resolution)^2^ and 3S3M (2.49 Å resolution)^3^ PDB structures, respectively. Protonation of all the amino acid residues for both proteins was predicted using y o ec e’s otein ep e o e t p. (30). Missing residues of the nucleosome were modelled based on PDB structure 1KX5 (21).

To predict the complexes between the trinucleosome and both the HIV-1 and PFV intasomes, the HADDOCK 2.4 web server was used (31). HADDOCK 2.4 was used for docking, applying a multi-step approach where solutions were grouped based on a 2.0 Å RMSD cutoff and ranked using HADDOCK scores. Active residues were selected based on a previous work (8). For the HIV-1 intasome, the selected residues were Y227, D229, R231, D232, W235, K236, and D253. For the human nucleosome the selected residues were R19, K20, V21, L22, and the 51-56 bp region. For the PFV intasome, active residues were selected based on the works of Maskell *et al* (24) and Wilson *et al* (26). For the nucleosome, these were K9 and R11 from the N-terminal of H2A and the C-terminal of H2B. (T119, S120, A121 and K122) as well as the target area of the DNA (36-41 pb). For the intasome, the residues selected as active were (P135, Q137, K168, A188, P239 and T240 of both chains).

## DATA AVAILABILITY

All data are available upon request.

## FUNDING

This work has been supported by the French ANRS research agency ECTZ115893, ECTZ227474, and FRM EQU202303016283.

## CONFLICT OF INTEREST DISCLOSURE

The authors declare no conflict of interest.

### AKNOWLEDGEMENTS

English editing has been performed by Nature Rearch Editing Service.

## SUP DATA

**S1.**
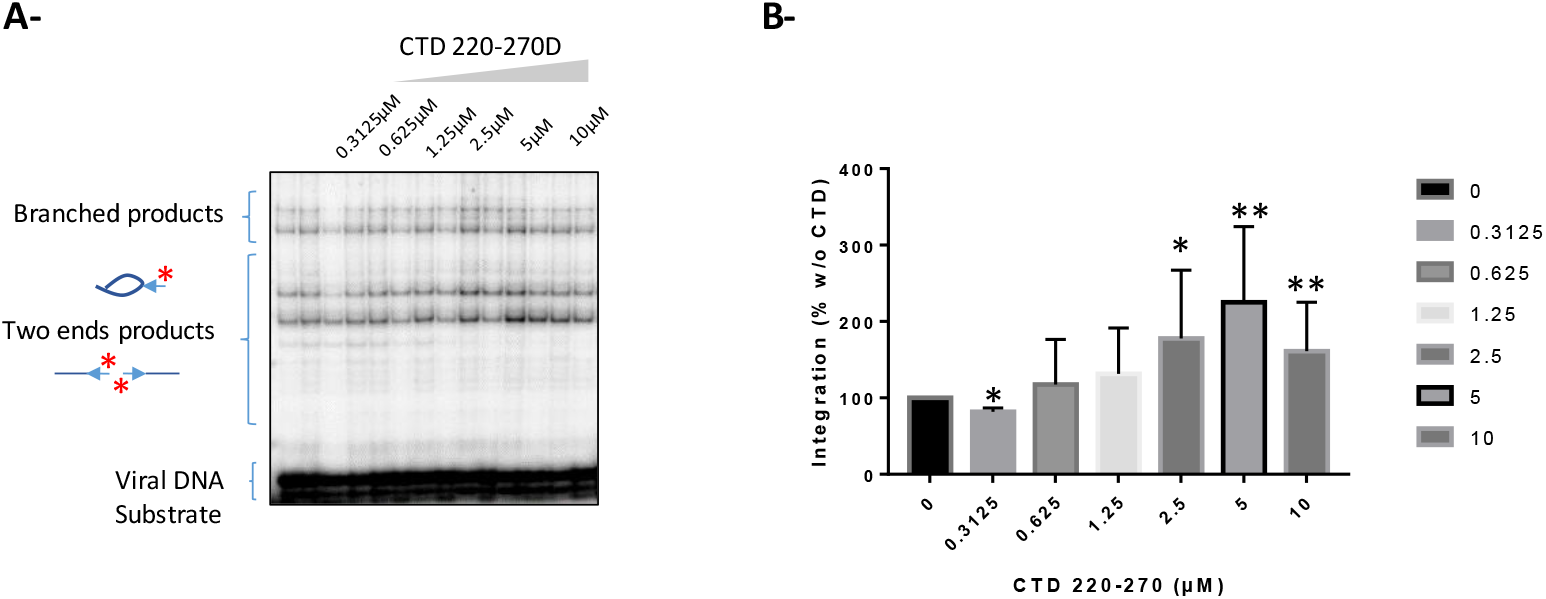
Effect of HIV-1 CTD on *in vitro* integration. Increasing concentrations of HIV-1 220-270 CTD were added to HIV-1 nucleosomal integration assay (**A**) and integration products were analyzed on 8% polyacrylamide gel. Integration products were quantified after autoradiography and data are reported as means of 3 independent experiments ±SD (**B**). T-student test was performed to determine the significance of the results: *p<0.01, **p<0.005.

**S2.**
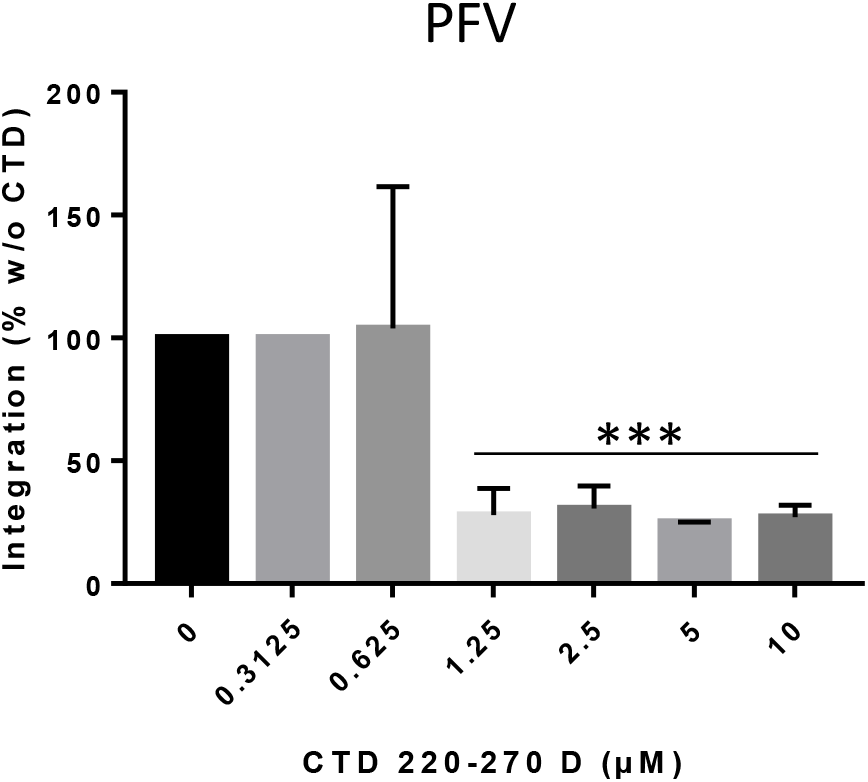
Effect of HIV-1 CTD on *in vitro* PFV integration. Increasing concentrations of HIV-1 220-270 CTD was added to PFV nucleosomal integration assay (**A**) and integration products were analyzed on 8% polyacrylamide gel. Integration products were quantified after autoradiography and data are reported as means of 3 independent experiments ±SD. T-student test was performed to determine the significance of the results: *p<0.01, **p<0.005. Integration assay was performed as in (**A**) and integration products were centrifugated before quantifying the radioactivity in the soluble fraction (**C**).

## Notes

### Competing Interest Statement

The authors have declared no competing interest.

### Summary of Updates

The first Author was miising in the previously submited MS. It has been fixed in this revised version.

## REFERENCES

1. Lesbats, P., Engelman, P.N. and Cherepanov, P. (2016) Retroviral DNA Integration. Chem Rev, 10.1021/acs.chemrev.6b00125.

2. Bedwell, P.J. and Engelman, P.N. (2021) Factors that mold the nuclear landscape of HIV-1 integration. Nucleic Acids Res., 49, 621–635.

3. Luger, P., Mader, P.W., Richmond, P.K., Sargent, P.F. and Richmond, P.J. (1997) Crystal structure of the nucleosome core particle at 2.8 A resolution. Nature, 389, 251–60.

4. Maertens, P.N., Engelman, P.N. and Cherepanov, P. (2021) Structure and function of retroviral integrase. Nat. Rev. Microbiol., 10.1038/s41579-021-00586-9.

5. Lesbats, P., Botbol, P., Chevereau, P., Vaillant, P., Calmels, P., Arneodo, P., Andreola, P.L., Lavigne, P. and Parissi, P. (2011) Functional Coupling between HIV-1 Integrase and the SWI/SNF Chromatin Remodeling Complex for Efficient in vitro Integration into Stable Nucleosomes. PLoS Pathog, 7, e1001280.

6. Benleulmi, P.S., Matysiak, P., Henriquez, P.R., Vaillant, P., Lesbats, P., Calmels, P., Naughtin, P., Leon, P., Skalka, P.M., Ruff, P., et al. (2015) Intasome architecture and chromatin density modulate retroviral integration into nucleosome. Retrovirology, 12, 13.

7. Maskell, P.P., Renault, P., Serrao, P., Lesbats, P., Matadeen, P., Hare, P., Lindemann, P., Engelman, P.N., Costa, P. and Cherepanov, P. (2015) Structural basis for retroviral integration into nucleosomes. Nature, 523, 366–369.

8. Benleulmi, P.S., Matysiak, P., Robert, P., Miskey, P., Mauro, P., Lapaillerie, P., Lesbats, P., Chaignepain, P., Henriquez, P.R., Calmels, P., et al. (2017) Modulation of the functional association between the HIV-1 intasome and the nucleosome by histone amino-terminal tails. Retrovirology, 14, 54.

9. Mauro, P., Lesbats, P., Lapaillerie, P., Chaignepain, P., Maillot, P., Oladosu, P., Robert, P., Fiorini, P., Kieffer, P., Bouaziz, P., et al. (2019) Human H4 tail stimulates HIV-1 integration through binding to the carboxy-terminal domain of integrase. Nucleic Acids Res., 10.1093/nar/gkz091.

10. Lapaillerie, P., Lelandais, P., Mauro, P., Lagadec, P., Tumiotto, P., Miskey, P., Ferran, P., Kuschner, P., Calmels, P., Métifiot, P., et al. (2021) Modulation of the intrinsic chromatin binding property of HIV-1 integrase by LEDGF/p75. Nucleic Acids Res., 10.1093/nar/gkab886.

11. Benleulmi, P.S., Matysiak, P., Robert, P., Miskey, P., Mauro, P., Lapaillerie, P., Lesbats, P., Chaignepain, P., Henriquez, P.R., Calmels, P., et al. (2017) Modulation of the functional association between the HIV-1 intasome and the nucleosome by histone amino-terminal tails. Retrovirology, 14, 54.

12. Li, P., Yang, P., Chen, P., Wang, P., Ghirlando, P., Dimitriadis, Emilios.K. and Craigie, P. (2024) HIV-1 integrase assembles multiple species of stable synaptic complex intasomes that are active for concerted DNA integration in vitro. J. Mol. Biol., 10.1016/j.jmb.2024.168557.

13. Passos, P.O., Li, P., Yang, P., Rebensburg, P.V., Ghirlando, P., Jeon, P., Shkriabai, P., Kvaratskhelia, P., Craigie, P. and Lyumkis, P. (2017) Cryo-EM structures and atomic model of the HIV-1 strand transfer complex intasome. Science, 355, 89–92.

14. Cook, P.J., Li, P., Berta, P., Badaoui, P., Ballandras-Colas, P., Nans, P., Kotecha, P., Rosta, P., Engelman, P.N. and Cherepanov, P. (2020) Structural basis of second-generation HIV integrase inhibitor action and viral resistance. Science, 367, 806–810.

15. Li, P., Li, P., Chen, P., Cui, P., Engelman, P.N. and Craigie, P. (2024) HIV-1 Intasomes Assembled with Excess Integrase C-Terminal Domain Protein Facilitate Structural Studies by Cryo-EM and Reveal the Role of the Integrase C-Terminal Tail in HIV-1 Integration. Viruses, 16, 1166.

16. Kimura, P., Saito, P., Hanada, P., Liu, P., Okabe, P., Kawashima, P.A., Yamatsugu, P. and Kanai, P. (2015) Supramolecular Ligands for Histone Tails by Employing a Multivalent Display of Trisulfonated Calix[4]arenes. ChemBioChem, 16, 2599–2604.

17. Mauro, P., Lapaillerie, P., Tumiotto, P., Charlier, P., Martins, P., Sousa, P.F., Métifiot, P., Weigel, P., Yamatsugu, P., Kanai, P., et al. (2023) Modulation of the functional interfaces between retroviral intasomes and the human nucleosome. mBio, 10.1128/mbio.01083-23.

18. Kalashnikova, P.A., Porter-Goff, P.E., Muthurajan, P.M., Luger, P. and Hansen, P.C. (2013) The role of the nucleosome acidic patch in modulating higher order chromatin structure. J. R. Soc. Interface, 10, 20121022.

19. Gamarra, P., Johnson, P.L., Trnka, P.J., Burlingame, P.L. and Narlikar, P.J. The nucleosomal acidic patch relieves auto-inhibition by the ISWI remodeler SNF2h. eLife, 7, e35322.

20. Matysiak, P., Lesbats, P., Mauro, P., Lapaillerie, P., Dupuy, P.-W., Lopez, P.P., Benleulmi, P.S., Calmels, P., Andreola, P.-L., Ruff, P., et al. (2017) Modulation of chromatin structure by the FACT histone chaperone complex regulates HIV-1 integration. Retrovirology, 14.

21. Davey, P.A., Sargent, P.F., Luger, P., Maeder, P.W. and Richmond, P.J. (2002) Solvent Mediated Interactions in the Structure of the Nucleosome Core Particle at 1.9Å Resolution. J. Mol. Biol., 319, 1097–1113.

22. Takizawa, P., Ho, P.-H., Tachiwana, P., Matsunami, P., Kobayashi, P., Suzuki, P., Arimura, P., Hori, P., Fukagawa, P., Ohi, P.D., et al. (2020) Cryo-EM Structures of Centromeric Tri-nucleosomes Containing a Central CENP-A Nucleosome. Structure, 28, 44–53.e4.

23. Hare, P., Smith, P.J., Metifiot, P., Jaxa-Chamiec, P., Pommier, P., Hughes, P.H. and Cherepanov, P. (Oct) Structural and Functional Analyses of the Second-Generation Integrase Strand Transfer Inhibitor Dolutegravir (S/GSK1349572). Mol Pharmacol, 80, 565–72.

24. Maskell, P.P., Renault, P., Serrao, P., Lesbats, P., Matadeen, P., Hare, P., Lindemann, P., Engelman, P.N., Costa, P. and Cherepanov, P. (2015) Structural basis for retroviral integration into nucleosomes. Nature, 523, 366–369.

25. Salentin, P., Schreiber, P., Haupt, P.J., Adasme, P.F. and Schroeder, P. (2015) PLIP: fully automated protein–ligand interaction profiler. Nucleic Acids Res., 43, W443–W447.

26. Wilson, P.D., Renault, P., Maskell, P.P., Ghoneim, P., Pye, P.E., Nans, P., Rueda, P.S., Cherepanov, P. and Costa, P. (2019) Retroviral integration into nucleosomes through DNA looping and sliding along the histone octamer. Nat. Commun., 10, 4189.

27. Koutná, P., Lux, V., Kouba, T., Škerloaá, P., Nováček, J., Srb, ., Hexnerosá, P., NŠoáchok, P., Kukačka, P., Novák, P., et al. (2023) Multivalency of nucleosome recognition by LEDGF. Nucleic Acids Res., 51, 10011–10025.

28. Lesbats, P., Botbol, P., Chevereau, P., Vaillant, P., Calmels, P., Arneodo, P., Andreola, P.L., Lavigne, P. and Parissi, P. (2011) Functional Coupling between HIV-1 Integrase and the SWI/SNF Chromatin Remodeling Complex for Efficient in vitro Integration into Stable Nucleosomes. PLoS Pathog, 7, e1001280.

29. Kimura, P., Saito, P., Hanada, P., Liu, P., Okabe, P., Kawashima, P.A., Yamatsugu, P. and Kanai, P. (2015) Supramolecular Ligands for Histone Tails by Employing a Multivalent Display of Trisulfonated Calix[4]arenes. ChemBioChem, 16, 2599–2604.

30. Torrens-Fontanals, P., Tourlas, P., Doerr, P. and De Fabritiis, P. (2024) PlayMolecule Viewer: A Toolkit for the Visualization of Molecules and Other Data. J. Chem. Inf. Model., 64, 584–589.

31. Honorato, P.V., Trellet, P.E., Jiménez-García, P., Schaarschmidt, P.J., Giulini, P., Reys, P., Koukos, P.I., Rodrigues, P.P.G.L.M., Karaca, P., Van Zundert, P.C.P., et al. (2024) The HADDOCK2.4 web server for integrative modeling of biomolecular complexes. Nat. Protoc., 19, 3219–3241.

